# CyAn: A MATLAB toolbox for image and data analysis of cyanobacteria

**DOI:** 10.1101/2020.07.28.225219

**Authors:** Jian Wei Tay, Jeffrey C. Cameron

## Abstract

Studies of photosynthetic cyanobacteria are often carried out by growing the bacteria in large volume cultures. However, this results in an uneven amount of light received by the cells due to cell-cell shading. Fluorescence microscopy solves this issue as cells can be grown in a single layer, eliminating shading issues. However, microscopy brings its own challenge as hundreds of cells are imaged at the same time, making it difficult to analyze the resulting data by hand. Here we present CyAn, a toolbox to process and analyze microscope images of cyanobacteria. The toolbox automates cyanobacteria identification in microscope images, tracks moving and dividing cells, and computes relevant biological data such as growth rates. As proof-of-principle, we demonstrate that wild type and mutants can be grown on the same slide and identified using the software, allowing a much more accurate probe into photosynthetic processes that are often masked due to cellular heterogeneity or those requiring precise light control to prevent cell-cell shading.

## Introduction

Cyanobacteria are ecologically important photosynthetic prokaryotes, often used as model organisms for research. As photoautotrophs, cyanobacteria convert CO_2_ and light energy into other biological products through photosynthesis. However, due to the requirement of light, photosynthetic research involving cyanobacteria in typical batch culture environments can be difficult. As the cells grow denser, light becomes attenuated through the culture making it difficult to control growth conditions (Berla et al. 2013). These self-shading effects can dramatically affect photosynthetic efficiency (Huisman and Weissing 1994; Kirk 1994; Hubble and Harper 2001; Passarge et al. 2006; Xiao et al. 2017).

A solution to this problem is to image cells using an optical fluorescence microscope. The microscope allows individual cells to be observed in vivo and in real time. An advantage of the microscope is that cells can be grown in a flat sheet, so each cell is exposed uniformly to the light. Furthermore, cyanobacteria are naturally fluorescent due to the presence of pigment-protein complexes in the internal thylakoid membrane system. In addition to providing signal for cell detection, the cellular fluorescence originating from photosystem I (PSII), photosystem II (PSII) and phycobilisome antenna proteins can also provide a quantitative and near instantaneous readout of the metabolic state of the cell (Campbell et al. 1998). In previous reports, fluorescence microscope imaging have been used to identify different photosynthetic species (Dijkman and Kromkamp 2006; Jin et al. 2018), as well as in studies on circadian gating in cyanobacterial cells (Dong et al. 2010; Yang et al. 2010).

However, a disadvantage of fluorescence microscopy is that extracting data from images is tedious and is often done a single frame over time. To solve this issue, we have developed CyAn, an object-oriented MATLAB toolbox written to extract and analyze data from microscope images of cyanobacteria cells. As a proof-of-principle demonstration, we show that the toolbox is useful for cyanobacteria photosynthesis research by filming wild type *Synechococcus* sp. PCC 7002 and a photosynthetic mutant lacking phycobilisomes (Δ*cpc*) at the same time. Since both strains are grown in under the same light and nutrient conditions, this technique eliminates uncertainty with the amount of incident light. The computational algorithms can identify individual cells, as well as track cell growth and division to form cell lineages. The computational analysis shows that this method can observe data at a range of length scales, from sub-cellular to colony-level.

## Method

### Strains and growth conditions

The species used in this work is the cyanobacterium *Synechococcus sp. PCC 7002* (hereafter PCC 7002). A mutant strain was created by genetic deletion of the *cpcB-F* operon (hereafter Δ*cpc*), which encodes for part of the phycobilisome antennae rods (Zhou et al. 1992). Liquid cultures of 25 mL in A+ media of both strains were grown from plates at 30°C, in air, and under white light provided by cool white fluorescent bulbs for 24 hours before imaging (∼200 μmol photons m^-2^ s^-1^). Just prior to imaging, each culture was diluted to an optical density of 0.05 OD at 730 nm. We found that at this density, cells were initially spread well apart so that the colonies do not merge during the experiment. 2 µL of the cells were then spotted on an agar pad (made with 0.5% agarose w/v A+ media), and the drop was allowed to dry completely (approximately 10 minutes) before placement in a glass bottom imaging dish (µ-Dish, Ibidi).

### Imaging setup

Imaging was carried out used a commercial mechanized microscope (Nikon TiE). Automated control of the microscope was performed by the supplied software (NIS-Elements AR version 5.11.00 64-bits). The sample stage of the microscope was enclosed by a cage incubator (Okolab) which provides both temperature and atmospheric control as necessary. For these experiments, the samples were grown in air at a temperature of 37 °C.

To ensure thermal stability, the temperature unit was turned on for at least an hour before imaging commenced. Furthermore, after the imaging dish was placed on the translational stage of the microscope, it was left for 30 minutes to allow the samples to reach thermal equilibrium before the microscope was focused. During this initial incubation period, the cells were illuminated with red light at an intensity of approximately 157 µmol photons m^-2^ s^-1^ to allow cells to acclimatize to the microscope environment. This process is important in maintaining focus throughout the experiment. After initial incubation, we used the built-in “perfect focus system” (PFS), which maintains the objective at the correct height from the bottom of the imaging dish, to focus the microscope. A total of 10 locations were then chosen in the NIS-Elements software for subsequent imaging.

All images were taken using a 100 x oil immersion objective (1.45 NA). We recorded images in three channels: chlorophyll fluorescence was imaged using the Cy5 filter (peak excitation of 645 nm, peak emission of 705 nm), phycobilisome fluorescence was imaged using the RFP filter (peak excitation of 555 nm, peak emission of 665 nm), and a brightfield image of the cells were acquired for validation purposes (Fig. 1). Note that the chlorophyll signal consist of both fluorescence emitted by photosystem II (PSII) pigments and the phycobilisomes in the wild type cells. In the Δ*cpc* strain, the chlorophyll fluorescence appears much dimmer and less uniform due to the lack of phycobilisome rods in this strain. The cells were imaged every 30 minutes for 12 hours, with a growth light of 299 µmol photons m^-2^ s^-1^. A representative video is available in the Supplementary Materials.

**Figure 1:**
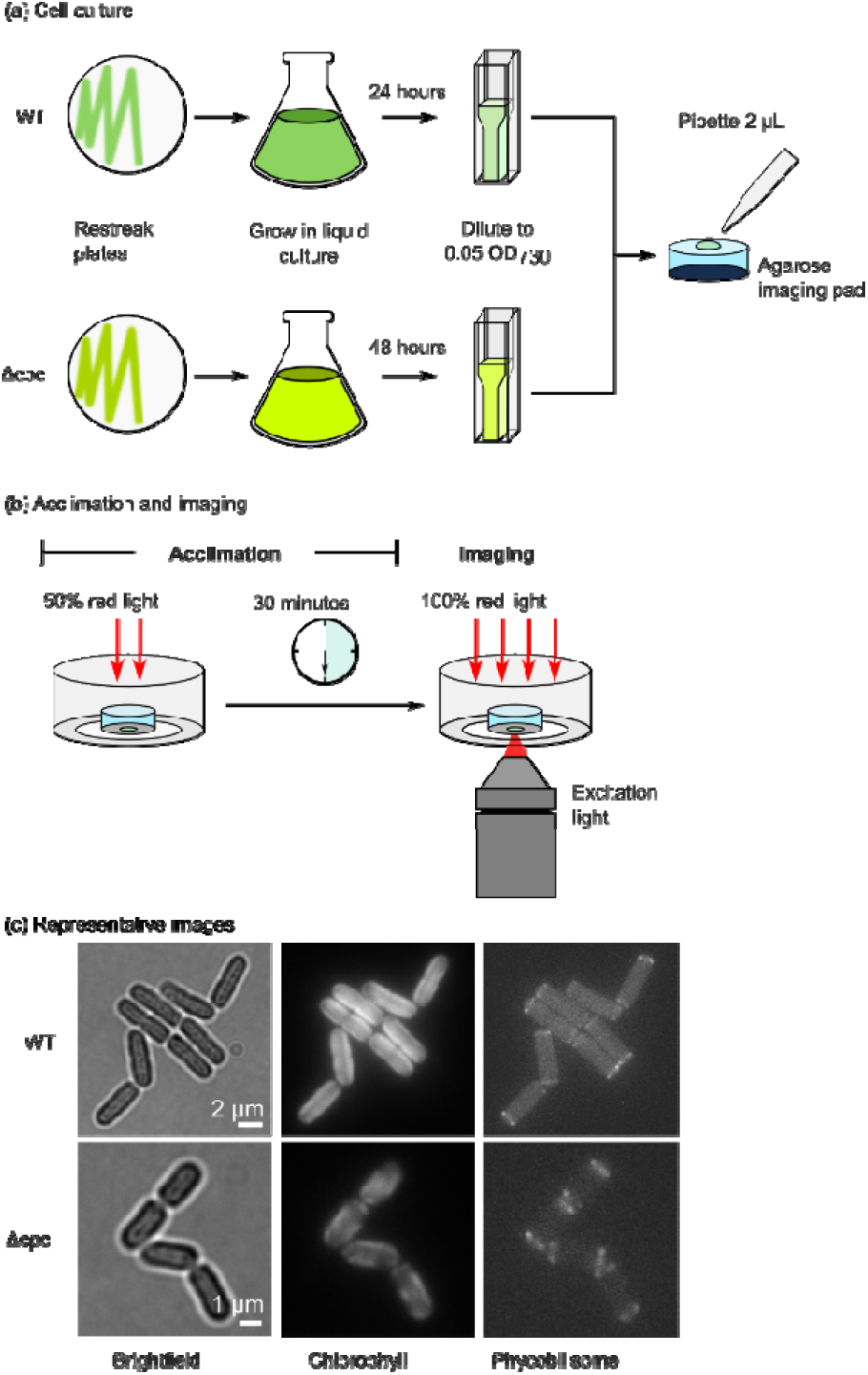
Experimental setup. (a) Wild type and Δcpc cells are cultured and plated on the same agarose imaging pad. (b) Cells are then placed on the microscope for 30 minutes to acclimate to the environment before imaging is carried out. (c) Representative images showing a typical wild type and Δcpc colony after 9 hours of growth. Note that the chlorophyll and phycobilisome fluorescence channels respectively have been normalized to the same values for both strains.

### Image analysis

To analyze the collected images, we developed a toolbox named CyAn. The toolbox was written in the MATLAB programming language and full code is available on the Gitlab platform (Tay). The workflow of the image analysis pipeline implemented in CyAn consists of four steps as shown in Fig. 2.

**Figure 2:**
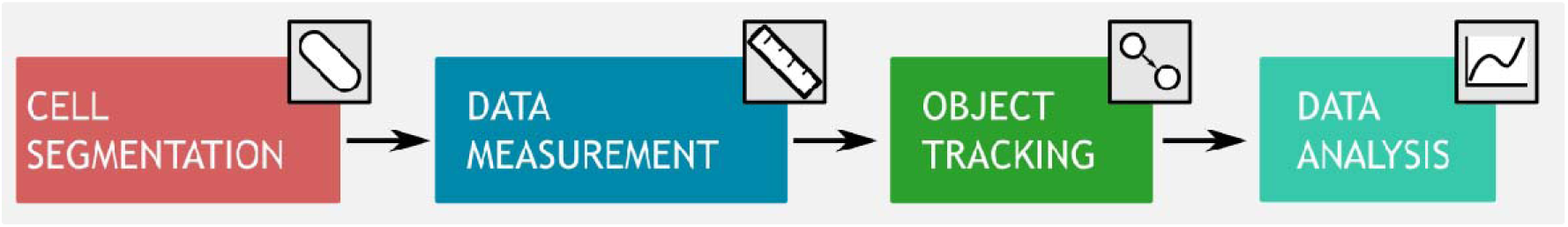
The image analysis pipeline taken by the CyAn software

In the first step, cells are identified in the image using a segmentation algorithm. The output of the segmentation process is a binary matrix also known as a mask. The mask has the same size as the image, with true pixels indicating pixels in the image belonging to a cell and false pixels indicating background. The mask is then used to measure cellular data, such as cell length, mean intensity, and position. The positional information is then used to identify the same cell through all frames of the movie to form a time series dataset of a cell, also known as a track. Finally, CyAn includes a data analysis class to analyze and visualize the resulting data. We describe the relevant algorithms for each step in this pipeline in the following sections.

### Cell segmentation

To identify the cells in the image, we used an adaptive threshold function (Bradley and Roth 2007) to generate an initial mask from the chlorophyll fluorescence image. We chose to use the fluorescence image as the cells are bright compared to the background. In contrast to the standard approach of choosing a single global intensity threshold, the adaptive threshold algorithm chooses different thresholds in different parts of the image, depending on local variation in the cell fluorescence. Once the initial mask was created, we removed pixels intersecting with the image border, as well as any objects less than 200 pixels in area. Finally, we applied the watershed function to separate cells that were clustered together. The results of the segmentation algorithm are shown in Fig. 3.

**Figure 3:**
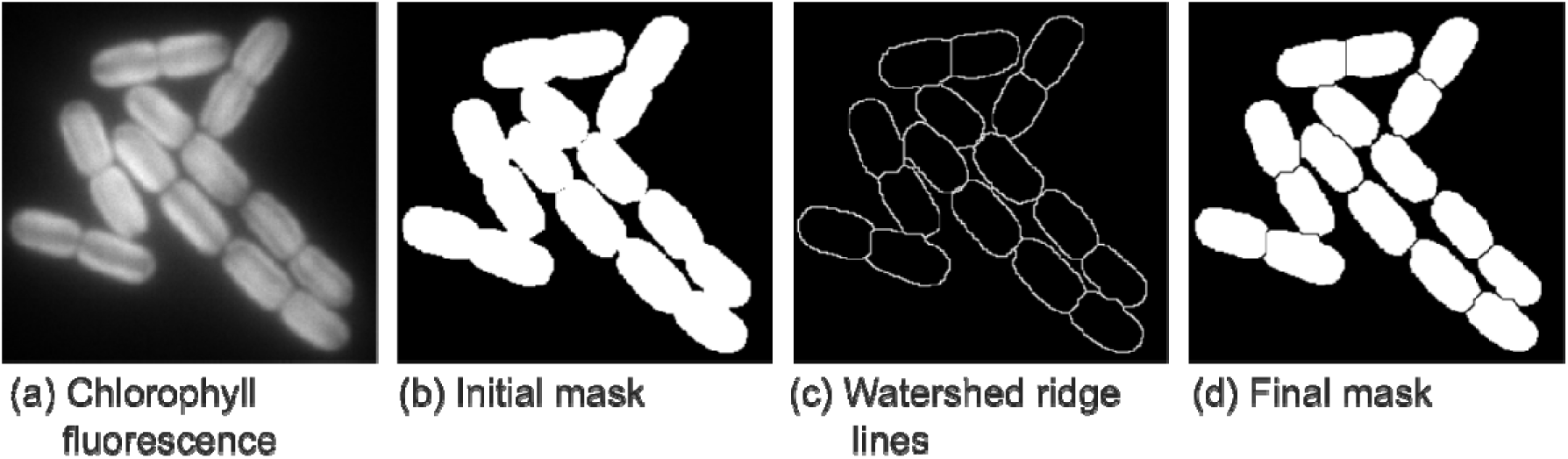
Representative figures from segmentation algorithm

### Measuring cell data

Cell data was then measured using the resulting mask. We measured the length and width of the cell by fitting an ellipse over the cell. For subsequent tracking of the cell, the index of each pixel belonging to a cell was also recorded.

### Tracking individual cells and cell division

Tracking is the process of following an individual cell over the entire movie. We used the linear assignment approach (Jaqaman et al. 2008) to perform tracking. In this approach, a score is calculated from the position of each object detected in the current frame to each object in the previous frame. This score, known as the linking cost, represents the probability that each pair of objects are the same individual; The lower the linking cost, the more likely it is that the objects are the same cell. In the original publication, the centroid position of each object was used to calculate the linking score. However, we found this method problematic as cyanobacteria cells are capsule shaped, which meant that the algorithm tended to link neighboring cells which were parallel to each other. We therefore used the ratio of overlapping pixels between frames instead.

To get a score for the degree of overlap, we use the ratio of the number of intersecting pixels with the number of union pixels, as shown in Fig. 4. Note that this overlap score ranges from 1 (cells are perfectly overlapping) to +Infinity (cells do not overlap at all). To block unlikely links (i.e. if cells only overlap by a few pixels), we set a threshold of 12. Higher linking scores were set to +Infinity indicating that the cells will not be linked by the algorithm. Scores for the alternative hypotheses that the newly detected object entered the field of view this frame (i.e. a new track should be created for this object) or that the previous object has left the field of view or become obscured by another cell (i.e. tracking should stop for that object) were also calculated using 1.05 * max(all linking scores). These scores were then assembled to form the cost matrix.

**Figure 4:**
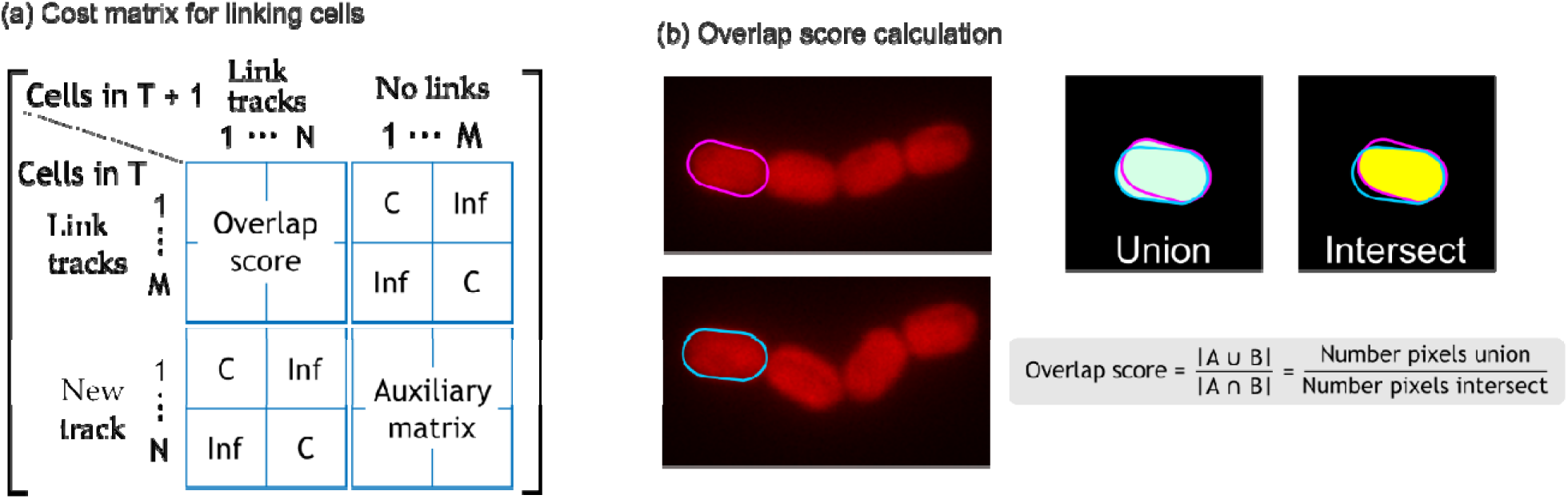
(a) Schematic of the linking matrix for linking cells between frames. The alternative linking score C is calculated as 1.05 * max(overlap scores). (b) The overlap score is calculated based on the number of overlapping pixels between two cell masks in subsequent frames.

A linear assignment algorithm was then used to assign a column to each row in the cost matrix. The assignment is made by minimizing the total cost of the assignments. In the original publication, the Kuhn-Munkres algorithm was used (Kuhn 1955; Munkres 1957). However, we found that this algorithm was slow when tracking large numbers of cells. Hence, we used the Jonker-Volgenant algorithm (Jonker and Volgenant 1987) instead. Internal testing showed that the Jonker-Volgenant algorithm solved the assignment problem ∼10x faster.

In addition to linking tracks, cell division was also identified by the linking algorithm. The probability of cell division was tested for when new tracks are created. Again, we used the overlap score to determine if division occurred, i.e. if two cells in the current frame overlapped at least 40% with the same mother cell in the previous frame.

By running the linking algorithm on subsequent frames, we were able to track individual cells over the course of the movie, as well as recording the resulting mother-daughter cell relationships. This time-series data was then stored as a structured array, which is a basic data type defined by MATLAB. We chose this format as it was a native data structure and would be easiest for future researchers to work with. The data for individual cells were stored as separate elements in the structured array, also called a track. The index of each track in the structure is unique. For each cell division, we kept track of the indices of the mother and resulting daughter cells, allowing us to analyze lineage, as well as to keep track of colonies in the movie.

### Image registration

Many movies experience a drift over time, caused by issues such as cycling of the heating system, drying of agarose pads, or changes in temperature of the microscope environment (Lee et al. 2012). Drift is problematic for tracking individual cells and must therefore be corrected. Here, we used the autofluorescence image of the cells as a fiduciary marker to register the images and correct for drifting.

Our system primarily experienced translational drift, i.e. the image shifted in x- and y-dimensions. To compute the drift correction, we calculated the cross-correlation between two frames of the movie of the chlorophyll fluorescence channel. The cross-correlation is given as where FFT is the (fast) Fourier transform, - is the inverse Fourier transform, and refers to the complex conjugate of the Fourier transform. The relative pixel shift is then given by the distance of the position corresponding to the maximum cross-correlation to the center of the image, as shown in **Fig. 5**. This drift correction was applied to the corresponding pixel positions before tracking.

**Figure 5:**
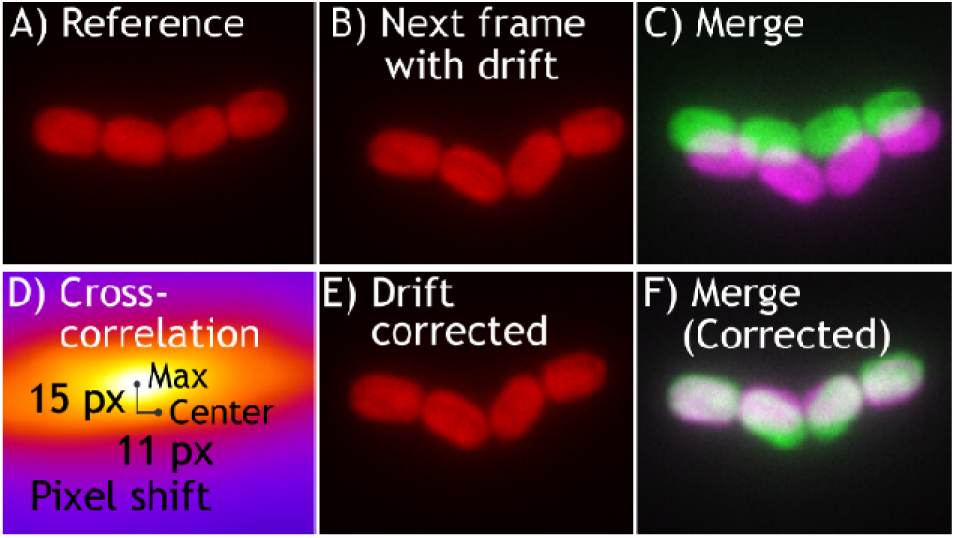
Process showing how image registration was carried out.

## Results

Using CyAn, we were able to record cell parameters such as cell length, in addition to chlorophyll and phycobilisome fluorescence intensity and cellular distribution. Furthermore, the tracking code records the unique indices assigned to mother and daughter cells, allowing analysis of traits within a single-cell derived lineage. In the following sections, we provide representative plots of useful parameters obtained.

### Identification of cell type

As the Δ*cpc* lacks full phycobilisomes, we can identify cell type from the RFP intensity measured for each cell. Fig. 6 shows representative results from this identification.

**Figure 6:**
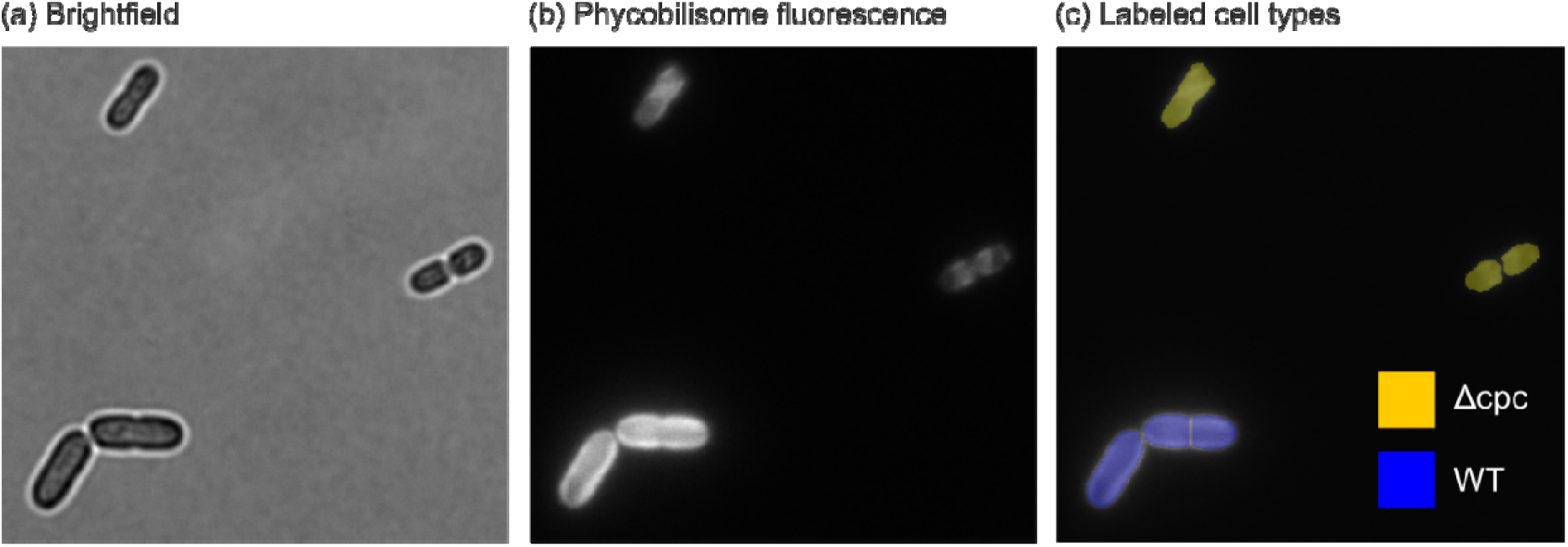
Cell type (WT or Δcpc) can be identified from the phycobilisome fluorescence intensity. (a) Shows the labeled cell types with Δcpc cells marked with a yellow overlay, and WT cells marked with blue. (b) The original phycobilisome fluorescence channel. (c) Brightfield image for confirmation (Δcpc cells are noticeably smaller than WT).

### Growth rates

The growth rate of each cell was found by fitting the measured cell length vs time to the equation

where *L*_*0*_ is the length of the cell at birth and k is the growth rate. The growth rate is related to the doubling time by

Fig. 7 shows a result for the fitting for a WT and a Δ*cpc* cell.

**Figure 7:**
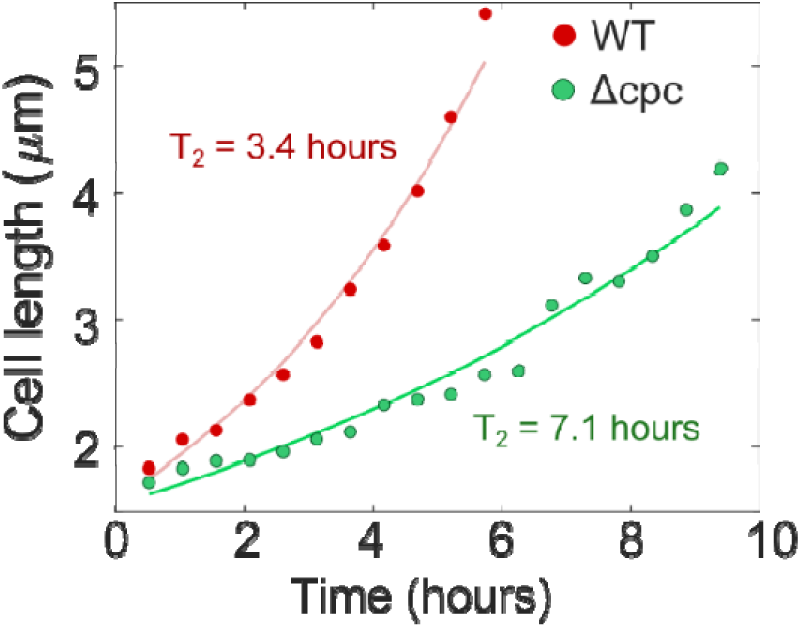
(a) Growth rates of WT (red) and Δcpc (green) cells. The original data points are shown as circles.

### Cell division and lineage tracing

As the tracking code records mother and daughter cell IDs, we can use this data to visualize the lineage of each cell. A function to plot the so-called lineage trees for a representative WT and Δ*cpc* lineage is shown in Fig. 8. This visualization can be a powerful tool in comparing the growth and eventual cell fate of different photosynthetic species.

**Figure 8:**
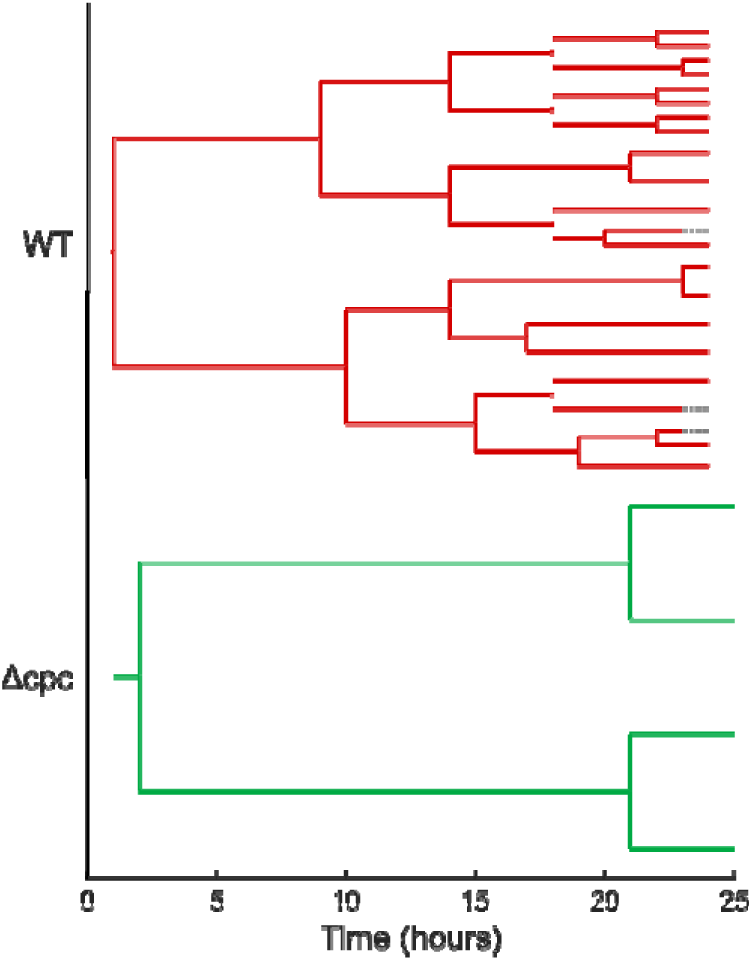
Lineage trees of representative cells. Note that the cells at the start of the movie exhibit a lag phase before starting to divide. The dotted lines in the wild type tree indicates when tracking was stopped for the cell due to segmentation issues as the colony grows in size.

## Conclusions

CyAn is an object-oriented MATLAB toolbox for analyzing microscope images of cyanobacteria. As a proof-of-principle, we collected data from actively growing cells of wild type PCC7002 and Δ*cpc*, a phycobilisome knockout mutant, co-cultured on the same agar pad. The single-cell phenotypes were then used to separate the individual strains computationally based on their fluorescent properties. This setup allows both strains to experience the exact same growth conditions. By tracking individual cellular lineages, population-level and emergent traits can be resolved at single-cell resolution. We believe that this technique will be valuable to the photosynthesis community as it allows a much more accurate probe into processes that are often masked due to cellular heterogeneity or those requiring precise light control to prevent cell-cell shading.

## Acknowledgements

We thank Dr. Kristin Moore for the Δ*cpc* strain. This study was financially supported in part by the U.S. Department of Energy (DOE) DE-SC0019306 (to J.C.C.).

## Code

The CyAn toolbox code is available at: https://biof-git.colorado.edu/cameron-lab/cyanobacteria-toolbox.

## Notes

### Competing Interest Statement

The authors have declared no competing interest.

